# Partial resistance to thyroid hormone-induced tachycardia and cardiac hypertrophy in mice lacking thyroid hormone receptor β

**DOI:** 10.1101/2023.11.21.567193

**Authors:** Riccardo Dore, Sarah Christine Sentis, Kornelia Johann, Nuria Lopez-Alcantara, Julia Resch, Lars Christian Moeller, Dagmar Fuehrer, Benedikt Obermayer, Robert Opitz, Jens Mittag

## Abstract

**Background:** Thyroid hormones regulate cardiac functions mainly via direct actions in the heart and binding to the thyroid hormone receptor (TR) isoforms α1 and β. While the role of the most abundantly expressed isoform, TRα1, is widely studied and well characterized, the role of TRβ in regulating heart functions is still poorly understood, primarily due to the accompanying elevation of circulating thyroid hormone in mice lacking TRβ (TRβ-KO). However, their hyperthyroidism is ameliorated at thermoneutrality, which allows studying the role of TRβ without this confounding factor.

**Methods:** Here we non-invasively monitored heart rate in TRβ-KO mice over several days using radiotelemetry at different housing temperatures (22°C and 30°C), and upon T3 administration in comparison to wildtype animals.

**Results:** TRβ-KO mice displayed normal average heart rate at both 22°C and 30°C with only minor changes in heart rate frequency distribution, which was confirmed by independent electrocardiogram recordings in freely-moving conscious mice. Parasympathetic nerve activity was, however, impaired in TRβ-KO mice at 22°C, and only partly rescued at 30°C. As expected, oral treatment with pharmacological doses of T3 at 30°C led to tachycardia in wildtypes, accompanied by broader heart rate frequency distribution and increased heart weight, while TRβ-KO mice showed blunted tachycardia, as well as resistance to changes in heart rate frequency distribution and heart weight. At the molecular level, these observations were paralleled by a blunted cardiac mRNA induction of several important genes, including the pacemaker channels *Hcn2* and *Hcn4*, as well as *Kcna7*.

**Conclusions:** The phenotyping of TRβ-KO mice conducted at thermoneutrality allows novel insights on the role of TRβ in cardiac functions in absence of the usual confounding hyperthyroidism. Even though TRβ is expressed at lower levels than TRα1 in the heart, our findings demonstrate an important role for this isoform in the cardiac response to thyroid hormones.

## Introduction

It has been long recognized that thyroid hormones (THs) tightly control cardiac activity, with tachycardia as one of the typical hallmarks of hyperthyroidism, and bradycardia observed in hypothyroidism (1, 2). THs, and specifically the active hormone 3,3’,5-triiodothyronine (T3), primarily act through regulating gene expression by the two nuclear receptors TRα1 and TRβ, which show a different pattern of expression throughout the body. For the heart, recent gene expression studies suggest that TRα1 is the major isoform; however, TRβ also accounts for ~30% of the total ligand-binding isoforms in the sinoatrial pacemaker cells and heart ventricles (3–5). Consequently, a number of studies have conclusively shown that mice carrying a mutation or deletion of TRα1 are bradycardic despite relatively normal circulating levels of THs (6–13). The role of TRβ in heart functions on the other hand is more controversial, as the knockout of TRβ leads to a hyperthyroid phenotype as a consequence of the disrupted negative feedback loop of the hypothalamus-pituitary-thyroid (HPT) axis - a confounding factor which renders the understanding of TRβ’s role in cardiac functions more difficult. Therefore, a number of studies observed a modest increase by 6-11% in heart rate in TRβ-KO mice, which is likely attributed to the elevated circulating THs levels and their action on the intact TRα1 (7, 9, 14). Interestingly, however, previous research reported an increase in heart rate of TRα1-KO mice upon T3, indicating that TRβ might also play an important role in the regulation of cardiac functions (12, 13).

In addition to the endocrine regulation, heart rate is also regulated by the autonomic nervous system (ANS), with the sympathetic branch increasing heart rate and the parasympathetic branch decreasing it (15). One major factor that greatly affects autonomic activity in both humans and rodents is the ambient temperature. In mice, it has been shown that the usual housing at room temperature (20-22°C) causes sympathetic activation with permanently elevated heart rate, and that even small changes in housing temperature (e.g. due to high housing density) can significantly affect heart rate in mice (16–19). Therefore, the conclusions from previous studies in rodents may have overestimated any cardiac phenotype due to the constant cold stress in animals housed at room temperature. To circumvent this issue, mice can be housed at thermoneutrality, a condition that is considered more translationally relevant (20–22). Even more importantly, as we have previously shown that housing at thermoneutrality strongly reduces the hyperthyroidism in TRβ-KO mice (23), this condition can eliminate confounding factors, thus allowing us to better study the role of TRβ in cardiac regulation.

In this study, we employed well-established radiotelemetry in conscious and freely moving TRβ-KO mice to long-term monitor heart rate and locomotor activity at room temperature (22°C) as well as at thermoneutrality (30°C). Furthermore, to better dissect the contribution of the sympathetic and parasympathetic nervous system in modulating cardiac activity, TRβ-KO mice were subjected to pharmacological denervation. Additionally, to conclusively clarify the role of the β isoform in thyroid hormone-induced tachycardia and cardiac hypertrophy, heart rate and weight were measured without and upon oral T3 treatment. Finally, the expression levels of several cardiac function-related markers (involved in pacemaking, repolarization and calcium handling) were quantified by quantitative real-time PCR following oral T3 treatment.

## Materials and Methods

### Animals Husbandry

Male TRβ-KO mice (24, 25) and control wildtypes were bred on the C57Bl/6NCrl background in the Gemeinsame Tierhaltung (GTH) of the University of Lübeck. Immediately after radiotelemetry transmitter implantation, animals were single housed in wire-topped, plastic cages (Techniplast, Italy) in a 12-hour light/dark cycle (lights on at 6:00 am), temperature-controlled (22 and 30 ± 1°C) air flow cabinet with *ad libitum* access to water and food (#1314, Altromin, Germany). Mice were ~5-month old at experimentś commencement. All animal experiments were carried out according to EU guideline regulations (210/63/EU) and approved by the MEKUN Schleswig-Holstein (Germany).

### In vivo Electrocardiogram

*In vivo* electrocardiograms (ECG) were recorded and analyzed using ECGenie system (Mouse Specifics, Inc., MA, USA). Mice were acclimated to an electrode-fitted platform (7 × 7.5 × 27 cm) for at least 10 minutes prior to data collection. On average, a number of 207 complexes per animal were analyzed.

### In vivo Radiotelemetry Recording

Implantable radio transmitters (Mini-Mitter Respironics, Bend, OR, USA) were used to determine heart rate and locomotor activity in conscious, undisturbed freely moving mice. Radio transmitters were implanted as described previously (11, 26–28) and animals recovered for 7 days prior to recording. Parameters were recorded every 30 s. For the long-term monitoring of heart rate and locomotor activity, samples were analyzed by calculating the 6-h average. To study the effect of housing temperatures and T3 on heart rate frequency distribution, the data were split into 20-bpm frequency bins and analyzed.

### Pharmacological Denervation

To study autonomic input and dissect sympathetic and parasympathetic contributions for control of heart rate, mice were intraperitoneally injected first with saline and, 45 minutes later, with scopolamine methyl bromide (0.1 mg/kg body weight; #S8502, Sigma-Aldrich, Germany) to block muscarinic receptors (PSNS). Finally, 45 minutes later, mice were injected with timolol maleate (1 mg/kg body weight; #T6394, Sigma-Aldrich, Germany) to block β-adrenergic receptors (SNS). Heart rate was constantly recorded through radiotelemetry. Pharmacological denervation experiments were conducted in the first half of inactive light phase. The median of each 45-min intervals and the differences in bpm were then calculated off-line (ΔPSNS: median [scopolamine] - median [saline]; ΔSNS: median [timolol] - median [scopolamine]). The intrinsic heart rate in bpm was recorded upon full receptors block (i.e. 45 minutes after timolol maleate administration).

### Pharmacological Treatment with Oral T3

Hyperthyroidism was induced at a housing temperature of 30 ± 1°C by treating the mice with 0.5 mg/L 3,3’,5-Triiodo-L-thyronine (#T6397, Sigma Aldrich, Germany) in 0.01% BSA and tap water for 12 days. Mice were provided every two days with freshly diluted solution and the water intake was monitored. Average daily dose of T3 was ~5µg/30g of BW, which was previously shown to result in a ~6- to 8-fold elevation in serum T3 and suppression in serum T4 (6, 29).

### Sacrifice, Organ Harvesting and RT-qPCR

Mice were euthanized using carbondioxide or isoflurane in combination with cervical dislocation. The hearts were quickly harvested, weighed, deep frozen in liquid nitrogen and stored at −80°C. Gene expression was performed as described previously (26, 30). The mRNA levels of target genes were normalized to those of *Cyclophilin D* and expressed in percentages to the wildtypes at 30°C. Sequences of gene-specific primers are provided in Supplementary Table S1.

### T3/T4 ELISA

After sampling, blood was centrifuged (1000 rpm for 10 minutes at 4°C) to obtain serum for subsequent total T3 (DNOV053; NovaTec Immundiagnostica GmbH, Germany) and T4 (EIA-1781; DRG Instruments GmbH, Germany) levels determination by following manufacture’s instructions.

### Re-analysis of Tabula muris senis single-cell data

Fully processed and annotated FACS single-cell data for mouse heart were downloaded from Figshare (https://figshare.com/ndownloader/files/23872838) and a dot plot was generated for selected genes using scanpy (v1.9.3; (31)).

### Statistical Analysis

All statistical analyses were performed using Excel (2016/2010/365, Version 2303) and GraphPad Prism 9.0 (GraphPad Software, US). Locomotor activity and heart rate were analyzed using two-way repeated measure (RM) analysis of variance (ANOVA) with genotype as between-subjects factor and time as within-subject factor or using unpaired Student’s *t*-test. Heart rate frequency distribution was analyzed using two-way RM-ANOVA with genotype as between-subjects factor and frequency bin as within-subject factor. Kurtosis and Skewness were analyzed using two-way RM-ANOVA with genotype as between-subjects factor and temperature as within-subject factor. Pharmacological denervation, intrinsic heart rate and ECG parameters were analyzed using unpaired Student’s *t*-test. The effect of T3 treatment was analyzed using two-way RM-ANOVA with genotype as between-subjects factor and time as within-subject factor. Normalized heart weight and gene expression were analyzed using two-way ANOVA with genotype and treatment as between-subjects. Total T3 and T4 serum levels were analyzed using either two-way ANOVA with genotype and temperature as between-subjects or unpaired Student’s *t*-test. *Post hoc* analysis was performed by using Sidak’s test. All data are expressed as mean ± standard error of the mean (SEM) and differences were considered statistically different at *p*<0.05. Further details can be found in Supplementary Table S2.

## Results

To gain more insights into the role of TRβ in the regulation of heart rate, the animals were non-invasively monitored using radiotelemetry at room temperature (22°C) and, subsequently, at thermoneutrality (30°C). Heart rate was not different between TRβ-KO and wildtype mice at neither temperature (Fig. 1A and B). As expected, a general reduction of heart rate by thermoneutrality was observed in both groups, indicating the shift from predominantly SNS to more PSNS control at 30°C. At 22°C, overall locomotor activity was reduced in TRβ-KO as compared with wildtype mice, but solely due to reduced activity during the dark active phase (−23%; Fig. 1C), comparable to what has been observed in a previous study of a hypothalamic TRβ knockdown (32). This difference in overall locomotor activity was no longer significant at 30°C (Fig. 1D). Total T3 and T4 serum levels of TRβ-KO mice were elevated by 75% compared to controls at 22°C, but not at 30°C (Suppl. Fig. 1A), as expected from previous studies (23). To further confirm the lack of tachycardia observed in the radiotelemetry experiments in spite of high TH levels at 22°C, we performed ECG recordings in another set of non-implanted mice. While we observed a significant shortening of QRS complexes duration in TRβ-KO mice, general heart rate remained unchanged between genotypes (Suppl. Fig. 1B and C).

**Figure 1:**
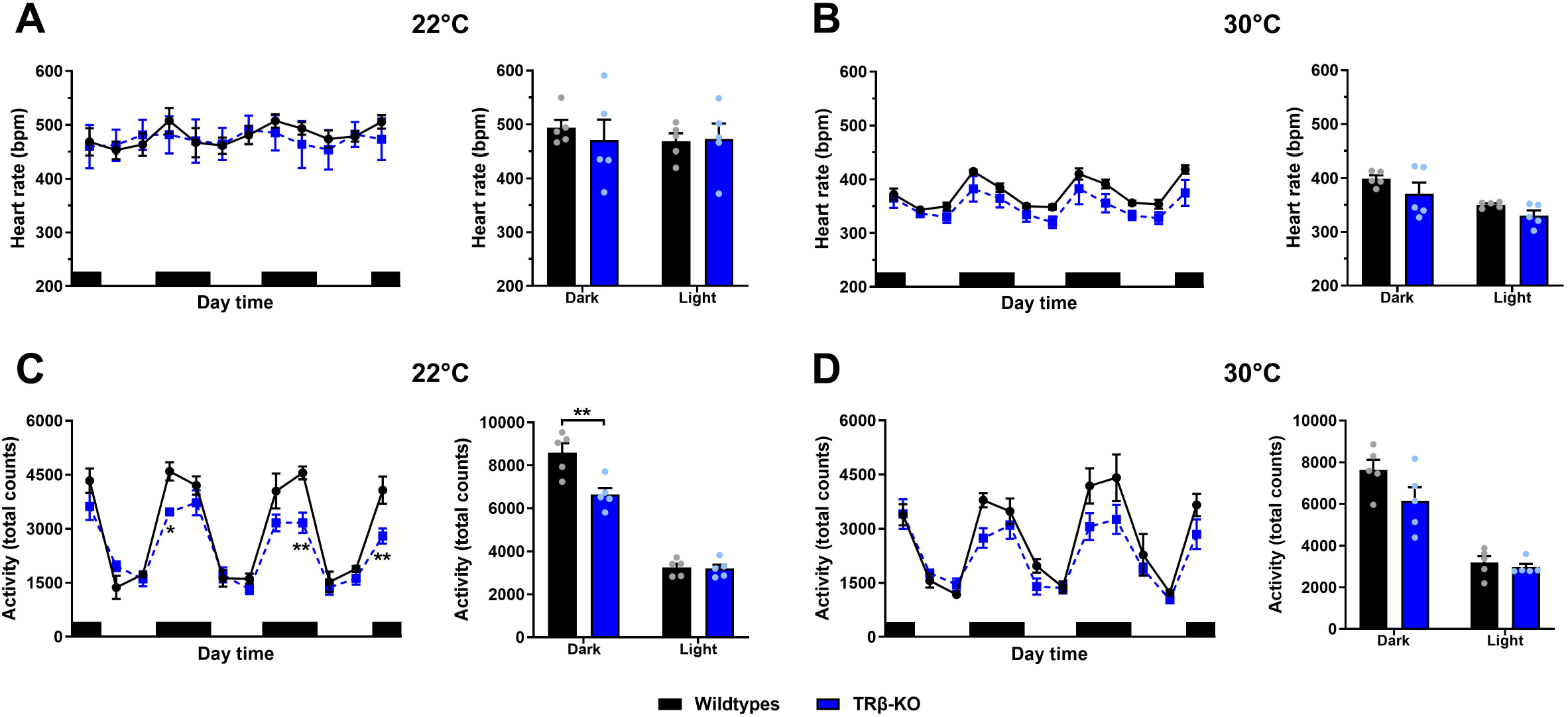
The effect of housing temperature on heart rate and locomotor activity in TRβ-KO mice. (A and B) Three-day radiotelemetry monitoring of heart rate frequency and (C and D) locomotor activity at 22°C and 30°C. (A and B) Average heart rate frequency and (C and D) locomotor activity during the dark active and light inactive phase at 22°C and 30°C. Black bars indicate dark active phases. Data represent mean ± SEM for TRβ-KO (blue; n=5) and wildtype controls (black; n=5). **P<0.01 (unpaired Studentśs *t* test).

To quantify heart rate variability, the 3-day-long heart rate monitoring was split into 20-bpm bins and the frequency distribution was analyzed. Even though the heart rate frequency distribution of TRβ-KO mice appeared to be broader than that of wildtypes, this finding was not significantly different (Fig. 2A). Again, when mice were housed at 30°C, there was a predominant PSNS control of heart rate as evidenced by a narrower distribution in both genotypes as compared with 22°C, leading to a visible temperature effect in kurtosis and skewness, which however failed to reach significance (Suppl. Fig. 2A). Interestingly, at 30°C, the heart rate frequency distribution was significantly narrower in TRβ-KO mice as compared to controls.

**Figure 2:**
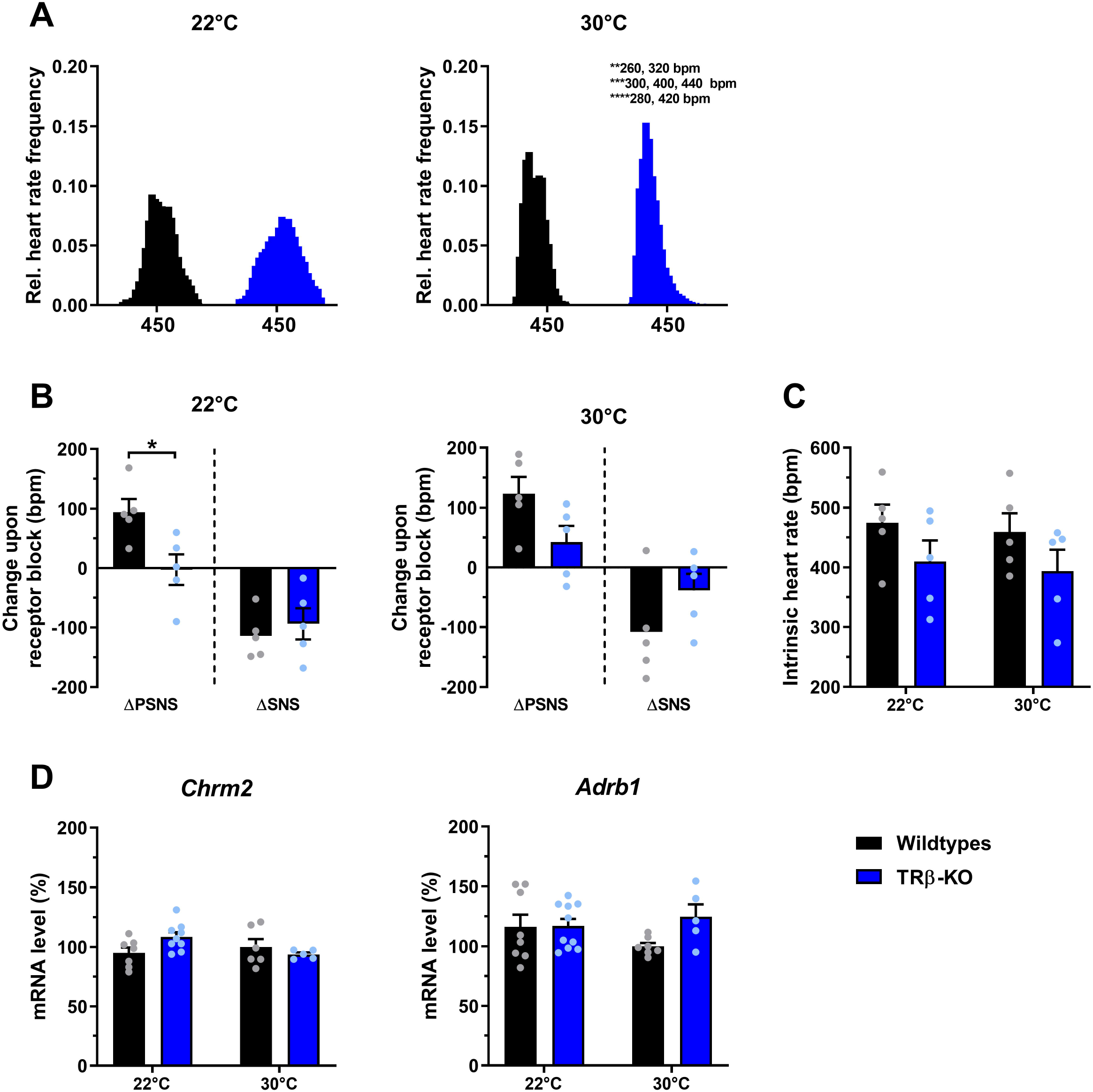
The effect of housing temperature on heart rate frequency distribution, pharmacological denervation and cardiac mRNA receptor levels in TRβ-KO mice. (A) Heart rate frequency distribution of three consecutive days at 22°C and 30°C. (B) Contributions of the parasympathetic (PSNS) or sympathetic nervous system (SNS) as determined by change in heart rate frequency upon pharmacological receptors blockade with scopolamine methyl bromide or timolol maleate at 22°C and 30°C. (C) The effect of housing temperature on intrinsic heart rate after full pharmacological receptors blockade, and (D) on cardiac expression of the muscarinic cholinergic receptor 2 (*Chrm2)* and adrenoceptor beta 1 *(Adrb1).* Data represent mean ± SEM for TRβ-KO (blue; n=5-10/group) and wildtype controls (black; n=5-8/group). *P<0.05, **P<0.01, ***P<0.001 and ****P<0.0001 (Sidak’s *post hoc* or unpaired Studentśs *t* test).

When we tested autonomic activity by using pharmacological blockade of PSNS and SNS *in vivo*, we observed impairment in the PSNS activity of TRβ-KO mice when housed at 22°C as indicated by a significantly reduced response to scopolamine methyl bromide. This was partially rescued at 30°C, as the residual 66% reduction was no longer significant, suggesting a beneficial effect of thermoneutrality on the PSNS activity in TRβ-KO (Fig. 2B). SNS activity did not differ between groups at any housing temperature as indicated by a comparable response to timolol maleate (Fig. 2B). Upon complete pharmacological autonomic receptor blockade, no change in intrinsic heart rate was observed in TRβ-KO mice at both temperatures (Fig. 2C). When we quantified mRNA expression levels of muscarinic receptor type 2 (*Chrm2*) and adrenoceptor type 1 (*Adrb1*) in the heart as a possible molecular mechanism underlying this pharmacological denervation response, we found no differences between groups at neither temperature (Fig. 2D).

Next, to induce tachycardia, we treated TRβ-KO and wildtype mice with T3 in the drinking water for 12 days at 30°C. As previously shown (6, 29), this pharmacological treatment led to a robust 5- to 7-fold elevation in serum levels of T3 and a parallel suppression in those of T4, with somewhat higher T3 levels in TRβ-KO mice (Suppl. Fig. 2B). As expected, while T3 treatment prominently increased heart rate in wildtype mice already after 2-3 days as compared to baseline, the effect was blunted in TRβ-KO mice resulting in a significantly reduced heart rate as compared to wildtypes (Fig. 3A). In addition, while the minimum heart rate reached during the light inactive phase was unaltered by T3 treatment in both groups, the maximum heart rate recorded in the dark active phase was significantly reduced in TRβ-KO as compared to controls (Suppl. Fig. 2C). On average, the delta heart rate measured between the dark and light phase was significantly smaller in TRβ-KO mice (Fig. 3B). As reported previously (6), the heart rate frequency distribution of wildtypes broadened upon T3, whereas that of TRβ-KO mice remained narrower as compared to the untreated condition and to wildtypes (Fig. 3C and Suppl. Fig. 2D and E). Heart weight was not different between groups at 30°C in untreated condition; however, T3 treatment resulted in a significant increase in the heart weight of wildtypes, which was not observed in TRβ-KO mice, resulting in a significant 14% reduction in heart weight compared to controls (Fig. 3D).

**Figure 3:**
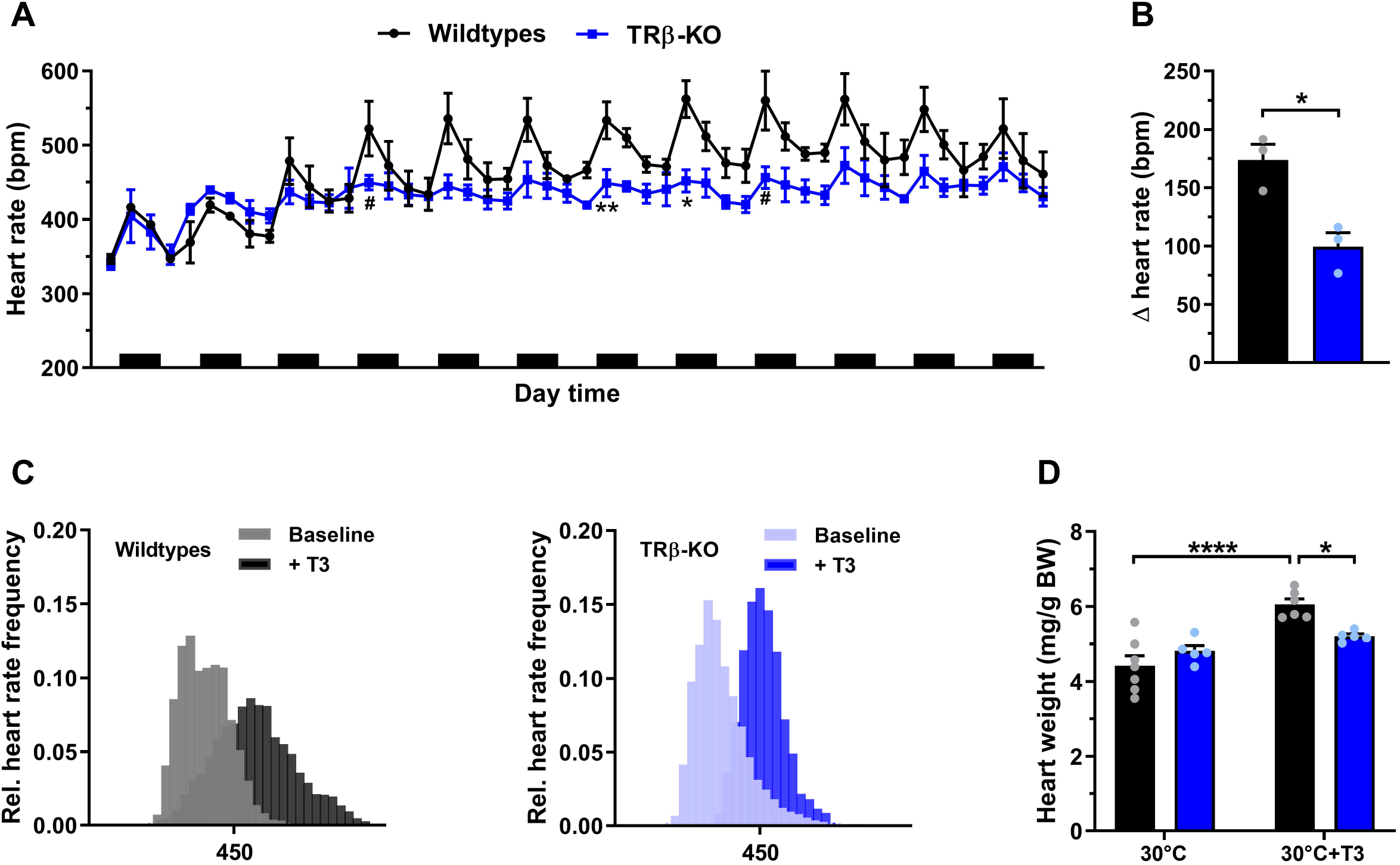
The effect of T3 treatment on heart rate, heart rate frequency distribution and heart weight in TRβ-KO mice at 30°C. (A) Radiotelemetry recordings of heart rate over 12 days of oral T3 treatment at 30°C. Black bars indicate dark active phases. (B) Average delta heart rate calculated as the difference between the maximum and minimum heart rate frequency recorded during dark and light phases, respectively. (C) Heart rate frequency distribution before and after T3 treatment at 30°C. (D) The effect of T3 treatment at 30°C on heart weight normalized by body weight. Data represent mean ± SEM for TRβ-KO (blue; n=3-5/group) and wildtype controls (black; n=3-7/group). ^#^P<0.1, *P<0.05, **P<0.01 and ****P<0.0001 (Sidak’s *post hoc* or unpaired Studentśs *t* test).

In line with the radiotelemetry data, the basal mRNA levels of the two pacemaker genes *Hcn2* and *Hcn4* were comparable to controls. Interestingly, while the expression of *Hcn2* was strongly induced by T3 in both genotypes, this was observed to a much lesser extent in TRβ-KO mice (+408% vs. 132%), resulting in significantly lower mRNA levels as compared to T3 treated wildtypes. Likewise, the expression of *Hcn4* was significantly induced only in wildtype animals (Fig. 4A). When we additionally quantified mRNA levels of other potassium channels implicated in cardiac repolarization, we found that similarly *Kcna7* was significantly induced by T3 treatment in wildtype but not in TRβ-KO mice. While the expression levels of *Kcnj3* and *Kcnq1* were significantly reduced by T3 treatment in both genotypes, those of *Kcnh2* were unresponsive to T3 and significantly lower in TRβ-KO mice as compared with wildtypes, suggesting that the proper transcription of this channel may require intact TRβ (Fig. 4A). When the ratio between *Myh6* and *Mhy7* was calculated as a measure of cardiac hypertrophy, the expected significant elevation in both groups was observed (Fig. 4A). Finally, we quantified the expression level of genes involved in calcium handling and cardiac contraction. While T3 treatment slightly but significantly lowered *Atp2a2* (Serca2) mRNA levels only in wildtype mice, those of *Pln*, an endogenous inhibitor of SERCA2 activity, were significantly decreased in both wildtype and TRβ-KO mice (Fig. 4A), resulting in a significant reduction in the ratio between *Pln* and *Atp2a2* in both genotypes (Suppl. Fig. 2F). The basal mRNA levels of *Ryr2* were significantly increased by 30% as compared to controls and interestingly, while T3 treatment enhanced *Ryr2* levels in wildtypes, it decreased them in TRβ-KO mice. However, there were no significant differences between wildtypes and TRβ-KO upon T3 in any of these calcium handling-related genes (Fig. 4A), suggesting that they are not involved in the observed partial resistance. Interestingly, when we measured *Dio2*, we observed generally low expression as expected, but a clear TRβ dependent acute regulation (Suppl. Fig. 2F). Finally, to better understand whether TRβ could be directly involved in the regulation of the tested genes, we reanalyzed published single-cell RNA sequencing data of adult mouse hearts to identify TRβ expressing cell types using the Tabula muris senis data set (33). These data showed expression of TRβ primarily in cardiomyocytes as well as lower TRβ expression in endocardial cells, smooth muscle cells and fibroblasts, which also expressed *Dio2* (Fig. 4B), thus supporting the possibility of a TRβ dependent regulation.

**Figure 4:**
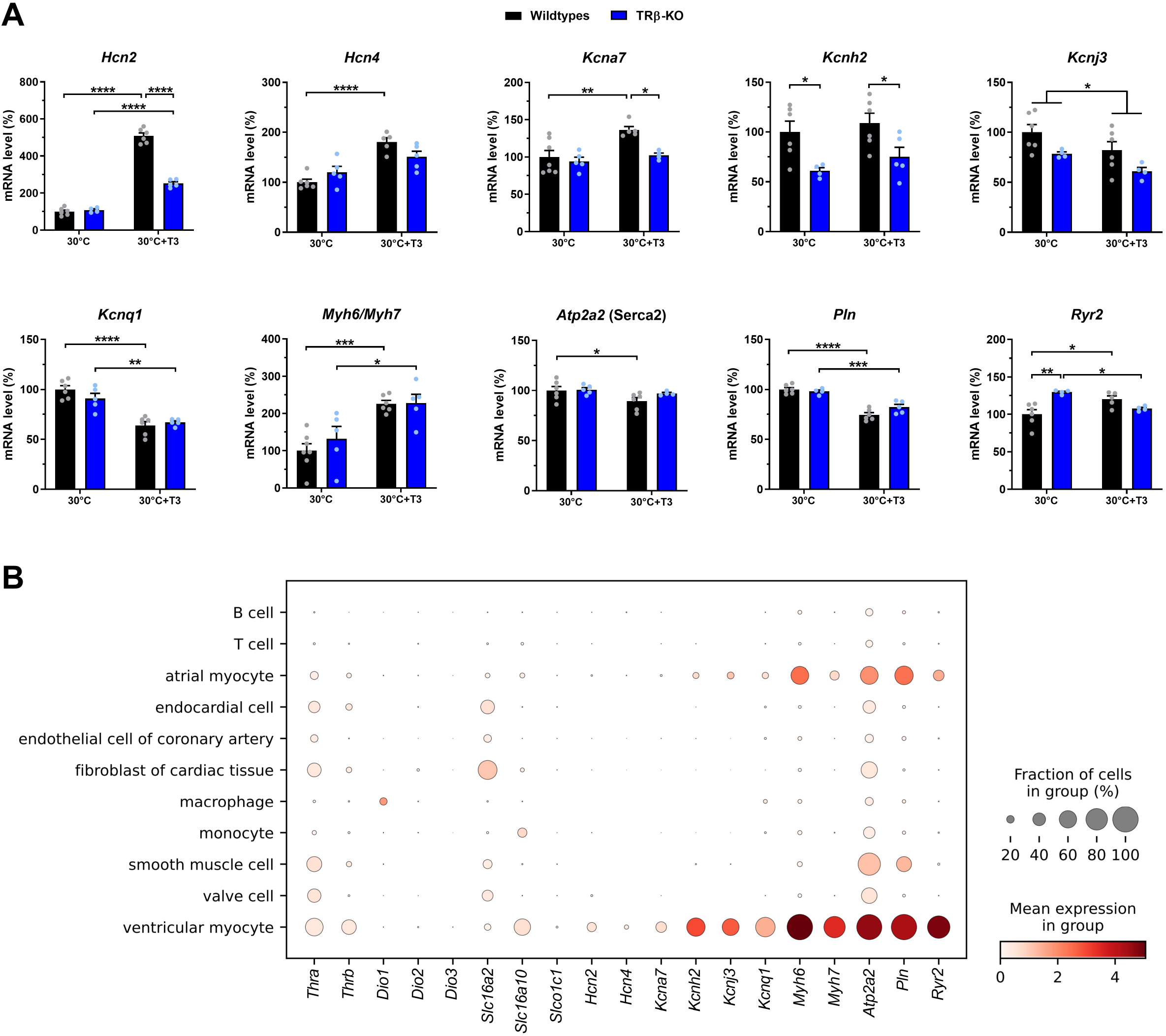
Cardiac gene expression with and without T3 treatment in TRβ-KO mice at 30°C. (A) Cardiac expression of genes involved in ‘pacemake’ (*Hcn2* and *Hcn4*, mediating the ‘funn’ potassium current), repolarization (*Kcna7*, mediating ultra-rapid potassium current; *Kcnh2*, mediating potassium rapid current; *Kcnj3*, mediating potassium acetylcholine-mediated current; *Kcnq1*, mediating potassium slow current), contraction and calcium handling (*Myh6* and *Myh7*, myosin heavy chain with fast and slow ATPase activity, respectively; *Atp2a2* (*Serca2*) *and Pln*, mediating calcium uptake into the sarcoplasmic reticulum and the inhibition of SERCA2 activity, respectively, and *Ryr2*, mediating the calcium extrusion from the sarcoplasmic reticulum. (B) Cell type-specific expression of selected genes in adult mouse hearts. Dot plot was generated based on published single cell RNA-seq data of the Tabula muris senis project. Dot size represents percentage of cells expressing the gene of interest in a given cell type and color denotes mean expression levels. Data represent mean ± SEM for TRβ-KO (blue; n=4-5/group) and wildtype controls (black; n=4-7/group). *P<0.05, **P<0.01, ***P<0.001 and ****P<0.0001 (Sidak’s *post hoc* test).

## Discussion

The main findings of the present study were: TRβ-KO mice showed (i) normal heart rate both at room temperature and thermoneutrality, (ii) moderately reduced locomotion and parasympathetic activity at room temperature, which were partially rescued at thermoneutrality; (iii) resistance to T3-induced tachycardia and cardiac hypertrophy, and (iv) altered expression of several cardiac activity-related genes, including *Hcn2*, *Kcna7* and *Ryr2*. Together, these findings suggest only a negligible role for TRβ under baseline conditions, but a more important contribution of the receptor in condition of systemic hyperthyroidism.

### TRβ is required for T3 induced cardiac hypertrophy

The induction of cardiac hypertrophy is one of the most classic effects triggered by THs (34). In addition to direct actions onto cardiomyocytes, THs-induced cardiac hypertrophy seems to be induced mainly, if not exclusively, *via* the modulation of the ANS (for review, see (35)). In fact, THs increase the expression/activity of β-adrenoceptors leading to a greater sensitivity of the heart to sympathetic stimulation and, eventually, to positive inotropic effects (36–38). Moreover, chronic administration of the β-adrenoceptors agonist isoproterenol increases heart weight (39, 40). Conversely, treatment with the β-adrenoceptors blocker propranolol inhibits T3-induced cardiac hypertrophy in parallel with increased heart rate (41). Corroborating these findings, while wildtype mice display cardiac hypertrophy after pharmacological treatment with either β-adrenoceptors agonist or T3, this effect was not observed in mice lacking all the β-adrenoceptors (42), indicating that T3-induced cardiac hypertrophy is to a large extend generated by the ANS. Consequently, Ortiga-Carvalho et al. (43) observed that while whole-body TRβ mutant mice developed cardiac hypertrophy upon T3 treatment, this effect was not displayed by animals with cardiac-specific TRβ mutation. In the present study, we found that control animals have a 36% increase in heart weight following T3 treatment, a clear indication of cardiac hypertrophy. This effect is not surprising considering that the pharmacological treatment employed here induced a 5- to 7-fold elevation in circulating T3 (6, 29) and that a significant increase in heart weight is detectable even upon lower doses of T3 (26). In contrast, TRβ-KO mice did not show similarly increased heart weight, suggesting that TRβ is required for the development of T3 induced cardiac hypertrophy at 30°C, an effect previously shown also at room temperature (43–45). While this effect cannot be simply attributed to acute differences in the *Myh6/Myh7* or *Pln/Atp2a2* (*Serca2*) ratio, as the levels were comparable to T3 treated wildtypes, it remains unclear whether this is the result of altered TRβ action in other tissues including the brain, or whether permanent alterations arising from the lack of TRβ during development could be involved.

### Lack of TRβ does not affect basal heart rate frequency at 22°C and 30°C

Previous studies have shown a modest increase in heart rate of TRβ-KO mice as a result of higher levels of THs; however, these experiments were all conducted in animals housed at room temperature, thus permanently exposed to minor cold stress (7, 9, 14). Here we show that the lack of TRβ has no effect on heart rate as evidenced by our *in vivo* radiotelemetry as well as ECG experiments at room temperature, indicating that the moderate ~75% elevation in THs levels observed at 22°C is not sufficient to induce tachycardia in mice. This observation is in complete accordance with our previous results showing a normal heart rate frequency profile in mice with similarly elevated TH levels (26).

A particular advantage of our present study is the phenotyping at thermoneutrality, a condition that better resembles that of humans, as animals are no longer cold-stressed and heart rate is predominantly under the control of the parasympathetic nervous system (21, 22). More importantly, we confirmed that housing at 30°C leads to a significant normalization of TH levels in TRβ-KO mice, allowing us to assess the contribution of the TRβ on heart functions without the confounding factors cold-stress and hyperthyroidism. Our *in vivo* data at thermoneutrality show normal heart rate in TRβ-KO mice together with comparable cardiac levels of the two pacemaker T3-target genes *Hcn2* and *Hcn4*. These results are in agreement with the notion that the expression of these two key pacemaker genes is mainly regulated by TRα1, and the previously observed increase in *Hcn2* and *Hcn4* in hyperthyroid tachycardic TRβ-KO mice (4, 46). When we induced a hyperthyroid state by T3 treatment for 12 days, TRβ-KO mice developed tachycardia but not to the same extent as the control animals, suggesting that TRβ may have an important role in allowing the heart to reach and sustain the maximum response/performance in a hyperthyroid state. The data are in agreement with a previous study showing a total failure of TRβ-KO mice to develop tachycardia upon T3 treatment (7); however, although the dose of T3 employed was comparable to that of our study, the treatment was restricted to only four days which may have been insufficient to generate any tachycardia in the somewhat resistant TRβ-KO mice. Most importantly, when TRα1-KO mice with intact TRβ signaling were treated with T3, a robust increase in heart rate was observed, strongly suggesting a contribution of TRβ in regulating heart activity upon T3 (12, 13). Our data of blunted tachycardia in T3 treated TRβ-KO mice match the molecular profile showing a reduced induction by T3 of the pacemaker gene *Hcn2* as well as the potassium channel *Kcna7*. It remains, however, to be determined whether this constitutes a developmental defect in TRβ-KO mice similar to that observed in TRα1 mutant mice (6, 30) or whether TRβ impairs cardiac adrenergic signaling by e.g. actions in other tissues, as this system has also been shown to be crucial for pacemaker gene induction (42).

### Altered PSNS activity and heart rate distribution in TRβ-KO mice

Since heart rate is also indirectly regulated through the modulation of the ANS, we aimed at dissecting the contributions of SNS and PSNS. Interestingly, while SNS activity was normal at both temperatures, TRβ-KO mice displayed reduced PSNS activity at 22°C, which was partially rescued by thermoneutrality. This improvement in PSNS activity seems not to be due to changes in muscarinic (*Chrm2*) and/or β-adrenergic (*Adrb1*) receptors as their gene expression remained unchanged. Furthermore, our observation of normal intrinsic heart rate indicates no cardiac defects caused by the lack of TRβ, suggesting that developmental and/or functional defects may reside in other structures (e.g. the hypothalamus). At present, it is difficult to establish whether the decrease in PSNS activity is related to the different T3 levels at 22°C and 30°C. While we observed that a 6-fold increase in T3 levels leads to decreased PSNS activity in mice housed at thermoneutrality (6), in another study we showed that mice housed at room temperature with moderately increased T3 levels similar to those of the TRβ-KO mice had normal PSNS activity (26), suggesting that only levels of T3 above a certain threshold may affect the PSNS. Given that previous studies showed normal SNS and PSNS activity in TRβ KO mice at room temperature (7), it seems likely that the effects of TRβ on PSNS activity are negligible unless they develop a strong hyperthyroid condition.

Another interesting result observed was the altered heart rate frequency distribution, with a generally broader distribution at 22°C and narrower at 30°C, indicative of a less stringent central control of heart rate at room temperature. In fact, while wildtypes mice show broader heart rate frequency distribution during T3 treatment as expected (6), TRβ-KO mice remained permanently narrow throughout the T3 treatment, suggesting that TRβ play an important role in adjusting heart rate stability in response to THs. This could be the result of developmental defects occurring in the central nervous system, as TRβ-KO mice show a ~40% reduction in the number of parvalbumin neurons in the anterior hypothalamic area, a pivotal brain area orchestrating autonomic, cardiovascular and stress response/functions (11, 47, 48). Whether this altered heart rate stability of the TRβ-KO mice is also detectable in response to other stimuli other than THs (e.g. stress, drugs, etc.) remains to be elucidated.

## Conclusions

Taken together, the present findings point towards a role of TRβ in regulating frequency distribution rather than average heart rate by a yet unknown mechanism possibly involving tissues other than the heart. More importantly, TRβ seems to be required for the full development of T3 induced tachycardia and hypertrophy; however, it remains unclear whether this is an acute effect or the consequence of the lack of TRβ actions during development.

## Acknowledgements

We thank the Gemeinsame Tierhaltung Lübeck for animal caretaking.

## Authorship Contribution Statement

Riccardo Dore, Sarah Christine Sentis and Jens Mittag conceptualized the study; Riccardo Dore, Sarah Christine Sentis, Kornelia Johann and Nuria Lopez-Alcantara performed the experiments; Julia Resch performed RT-qPCR; Riccardo Dore, Benedikt Obermayer, Robert Opitz and Jens Mittag analyzed data; Riccardo Dore, Lars Christian Moeller, Dagmar Führer, Benedikt Obermayer, Robert Opitz and Jens Mittag interpreted data; Riccardo Dore and Jens Mittag drafted the manuscript. All authors provided critical revision of the manuscript and approved its final version for publication.

