## Supplemental materials for "Partial resistance to thyroid hormone-induced tachycardia and cardiac hypertrophy in mice lacking thyroid hormone receptor β"

**Supplementary Figure 1**

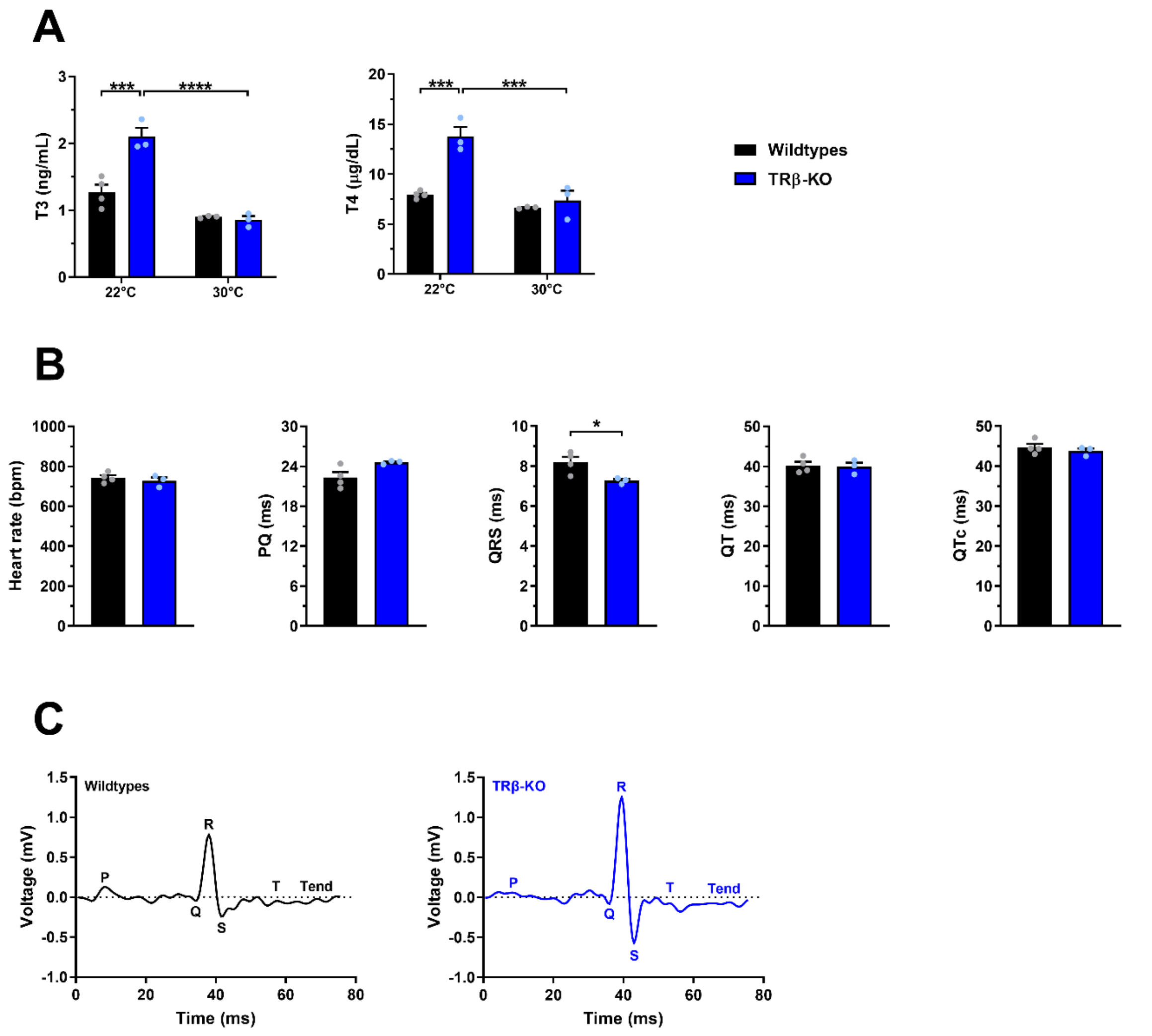


**Supplementary Figure 1: TH levels at different housing temperatures and ECG parameters**

(A) Total T3 and T4 serum levels in wildtype and TRβ knockout animals housed at 22°C and 30°C (related to Fig 1A-D). (B) Heart rate as well as other ECG parameters in these mice (related to Fig 2A). (C) Representative QRS complexes of wildtype and TRβ-KO mice. Data represent mean ± SEM. *P<0.05, ***P<0.001 and ****P<0.0001 (Sidak´s *post hoc* test or unpaired Student's *t* test).

### Supplementary Figure 2

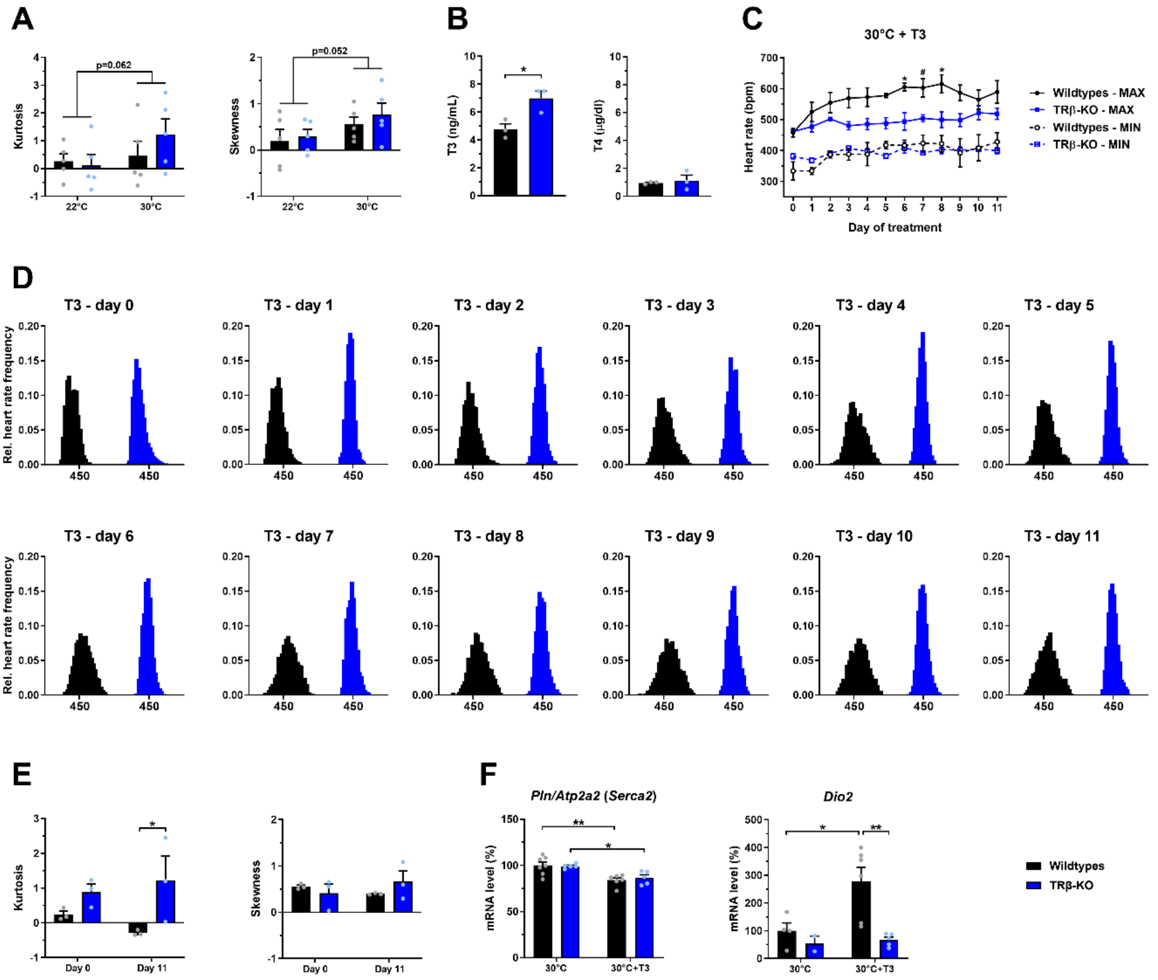


**Supplementary Figure 2: Effect of T3 treatment on heart functions, TH levels and gene expression**

(A) The effect of housing temperature on heart rate frequency distribution as measured by kurtosis and skewness (related to Fig. 2A). (B) Total T3 and T4 serum levels after T3 treatment for 12 days at 30°C (related to Fig. 3A). (C) Daily maximum and minimum heart rate frequency recorded in the dark and light phases, respectively, during T3 treatment for 12 days at 30°C (related to Fig. 3A). (D) Day-by-day development of heart rate frequency distribution over 12 days of T3 treatment at 30°C. (E) The effect to T3 treatment for 12 days at 30°C on heart rate frequency distribution as measured by kurtosis and skewness. (F) Cardiac *Pln/Atp2a2* (*Serca2*) ratio and expression of *Dio2*. Data represent mean ± SEM or fractions of 20-bpm bins. $^{\#}$<0.1, *P<0.05 and **P<0.01 (Sidak´s *post hoc* test or unpaired Students´s *t* test)

**Supplementary Table 1: Gene-specific primer sequences**

| Gene | Forwards (5´-3´) | Reverse (5´-3´) |
|---|---|---|
| *Adrb1* | CTCATCGTGGTGGGTAACGTG | ACACACAGCACATCTACCGAA |
| *Atp2a2* (*Serca2*) | TCCGCTACCTCATCTCATCC | CAGGTCTGGAGGATTGAACC |
| *Chrm2* | CAAGATCCAGAATGGCAAGG | GACAGACGTGGAGTCATTGG |
| *Cyclophilin D* | TCACAACAGTTCCGACTCCTC | ACCTCTACATTTTCAAGCGTCC |
| *Dio2* | ATGGGACTCCTCAGCGTAGAC | ACTCTCCGCGAGTGGACTT |
| *Hcn2* | TCCGCACCGGCAAAGTTA | CCGGGATGGATGACACGAAG |
| *Hcn4* | ACCCGCAGAGGATCAAGATGA | AATGCGAGTCTCCACTATAAGGA |
| *Kcna7* | CCATCCTAAGGGTCATCCGAT | TGAGGAGACCTAGCTCACGC |
| *Kcnh2* | GTGCTGCCTGAGTATAAGCTG | CCGAGTACGGTGTGAAGACT |
| *Kcnj3* | GGGGACGATTACCAGGTAGTG | CGCTGCCGTTTCTTCTTGG |
| *Kcnq1* | ACCGTCTTCCTCATTGTTCTGG | GACAATCTCCATCCAGAAGAGG |
| *Myh6* | GCCCTTTGACATTCGCACTG | GGTTTCAGCAATGACCTTGCC |
| *Myh7* | CCTGCGGAAGTCTGAGAAGG | CTCGGGACACGATCTTGGC |
| *Pln* | ACTGTGACGATCACCGAAGC | TTCCATTATGCCAGGAAGG |
| *Ryr2* | ACGGCGACCATCCACAAAG | CGGGGGAACATTCTTGGAATT |

**Supplementary Table 2: Statistical Details for all Figures**

| | | | 2-way ANOVA | | |
|---|---|---|---|---|---|
| | | | Interaction | Genotype | Time |
| Figure 1A | 22°C | | F(11, 88)=0.92<br>P=0.5236 | F(1, 8)=0.05<br>P=0.8299 | F(11, 88)=2.12<br>P=0.0268 |
| | | | Unpaired Student´s t-test | | |
| | | | Df | t | P value |
| Figure 1A | 22°C | Dark | 8 | 0.562 | 0.5894 |
| | | Light | 8 | 0.132 | 0.8981 |
| | | | 2-way ANOVA | | |
| | | | Interaction | Genotype | Time |
| Figure 1B | 30°C | | F(11, 88)=0.85<br>P=0.5888 | F(1, 8)=2.56<br>P=0.1481 | F(11, 88)=17.90<br>P<0.0001 |
| | | | Unpaired Student´s t-test | | |
| | | | Df | t | P value |
| Figure 1B | 30°C | Dark | 8 | 1.273 | 0.2389 |
| | | Light | 8 | 2.045 | 0.0751 |
| | | | 2-way ANOVA | | |
| | | | Interaction | Genotype | Time |
| Figure 1C | 22°C | | F(11, 88)=2.89<br>P=0.0027 | F(1, 8)=13.37<br>P=0.0064 | F(11, 88)=44.44<br>P<0.0001 |
| | | | Unpaired Student´s t-test | | |
| | | | Df | t | P value |
| Figure 1C | 22°C | Dark | 8 | 3.665 | 0.0064 |
| | | Light | 8 | 0.193 | 0.8517 |
| | | | 2-way ANOVA | | |
| | | | Interaction | Genotype | Time |
| Figure 1D | 30°C | | F(11, 88)=1.66<br>P=0.0964 | F(1, 8)=3.15<br>P=0.1140 | F(11, 88)=26.89<br>P<0.0001 |
| | | | Unpaired Student´s t-test | | |
| | | | Df | t | P value |
| Figure 1D | 30°C | Dark | 8 | 1.817 | 0.1067 |
| | | Light | 8 | 0.691 | 0.5089 |

| | | | 2-way ANOVA | | |
|---|---|---|---|---|---|
| | | | Interaction | Genotype | Bin |
| Figure 2A | 22°C | | F(35, 280)=0.79<br>P=0.7970 | F(1, 8)=0.00<br>P>0.9999 | F(35, 280)=15.49<br>P<0.0001 |
| | 30°C | | F(35, 280)=5.57<br>P<0.0001 | F(1, 8)=4.00<br>P=0.0805 | F(35, 280)=82.14<br>P<0.0001 |
| | | | Unpaired Student´s t-test | | |
| | | | Df | t | P value |
| Figure 2B | 22°C | ΔPSNS | 8 | 2.882 | 0.0205 |
| | | ΔSNS | 8 | 0.632 | 0.5448 |
| | 30°C | ΔPSNS | 8 | 2.074 | 0.0718 |
| | | ΔSNS | 8 | 1.507 | 0.1702 |

| | | | 2-way ANOVA | | |
|---|---|---|---|---|---|
| | | | Interaction | Genotype | Temperature |
| Figure 2C | | | F(1, 8)=0.00<br>P=0.9795 | F(1, 8)=2.46<br>P=0.1552 | F(1, 8)=0.50<br>P=0.4975 |
| Figure 2D | | *Chrm2* | F(1, 23)=4.32<br>P=0.0490 | F(1, 23)=0.58<br>P=0.4546 | F(1, 23)=1.06<br>P=0.3128 |
| | | *Adrb1* | F(1, 26)=2.30<br>P=0.1412 | F(1, 26)=2.60<br>P=0.1192 | F(1, 26)=0.28<br>P=0.5996 |

| | | | 2-way ANOVA | | |
|---|---|---|---|---|---|
| | | | Interaction | Genotype | Time |
| Figure 3A | 30°C | | F(47, 188)=3.05<br>P<0.0001 | F(1, 4)=3.39<br>P=0.1392 | F(47, 188)=13.62<br>P<0.0001 |
| | | | Unpaired Student´s t-test | | |
| | | | Df | t | P value |
| Figure 3B | 30°C | | 4 | 4.132 | 0.0145 |
| | | | 2-way ANOVA | | |
| | | | Interaction | Genotype | Treatment |
| Figure 3D | 30°C | | F(1, 19)=9.94<br>P=0.0052 | F(1, 19)=1.32<br>P=0.2652 | F(1, 19)=26.12<br>P<0.0001 |

| | | | 2-way ANOVA | | |
|---|---|---|---|---|---|
| | | | Interaction | Genotype | Treatment |
| Figure 4A | 30°C | *Hcn2* | F(1, 17)=126.70<br>P<0.0001 | F(1, 17)=111.70<br>P<0.0001 | F(1, 17)=549.60<br>P<0.0001 |
| | | *Hcn4* | F(1, 17)=6.95<br>P=0.0173 | F(1, 17)=0.26<br>P=0.6157 | F(1, 17)=34.87<br>P<0.0001 |
| | | *Kcna7* | F(1, 17)=4.00<br>P=0.0617 | F(1, 17)=8.03<br>P=0.0114 | F(1, 17)=10.02<br>P=0.0057 |
| | | *Kcnh2* | F(1, 17)=0.07<br>P=0.7872 | F(1, 17)=13.69<br>P=0.0018 | F(1, 17)=1.35<br>P=0.2604 |
| | | *Kcnj3* | F(1, 16)=0.00<br>P=0.9751 | F(1, 16)=8.52<br>P=0.0100 | F(1, 16)=5.79<br>P=0.0285 |
| | | *Kcnq1* | F(1, 18)=2.56<br>P=0.1268 | F(1, 18)=0.64<br>P=0.4341 | F(1, 18)=60.71<br>P<0.0001 |
| | | *Myh6/Myh7* | F(1, 19)=0.50<br>P=0.4890 | F(1, 19)=0.63<br>P=0.4380 | F(1, 19)=26.26<br>P<0.0001 |
| | | *Atp2a2*<br>(*Serca2*) | F(1, 17)=1.03<br>P=0.3237 | F(1, 17)=1.61<br>P=0.2215 | F(1, 17)=4.84<br>P=0.0419 |
| | | *Pln* | F(1, 17)=4.90<br>P=0.0408 | F(1, 17)=1.60<br>P=0.2228 | F(1, 17)=85.19<br>P<0.0001 |
| | | *Ryr2* | F(1, 15)=19.61<br>P=0.0005 | F(1, 15)=3.19<br>P=0.0942 | F(1, 15)=0.04<br>P=0.8419 |

<table>
<tr><td colspan="3" rowspan="2"></td><td colspan="3">2-way ANOVA</td></tr>
<tr><td>Interaction</td><td>Genotype</td><td>Temperature</td></tr>
<tr><td rowspan="2">Supp Fig 1A</td><td>T3</td><td></td><td>F(1, 9)=20.38<br>P=0.0015</td><td>F(1, 9)=16.36<br>P=0.0029</td><td>F(1, 9)=70.22<br>P<0.0001</td></tr>
<tr><td>T4</td><td></td><td>F(1, 9)=16.40<br>P=0.0029</td><td>F(1, 9)=26.96<br>P=0.0006</td><td>F(1, 9)=36.96<br>P=0.0002</td></tr>
<tr><td colspan="3" rowspan="2"></td><td colspan="3">Unpaired Student´s t-test</td></tr>
<tr><td>Df</td><td>t</td><td>P value</td></tr>
<tr><td rowspan="5">Supp Fig 1A</td><td rowspan="5">22°C</td><td>Heart rate</td><td>5</td><td>0.7577</td><td>0.4828</td></tr>
<tr><td>PQ</td><td>5</td><td>2.4020</td><td>0.0615</td></tr>
<tr><td>QRS</td><td>5</td><td>2.9020</td><td>0.0337</td></tr>
<tr><td>QT</td><td>5</td><td>0.2156</td><td>0.8378</td></tr>
<tr><td>QTc</td><td>5</td><td>0.8119</td><td>0.4537</td></tr>
</table>

<table>
<tr><td colspan="3" rowspan="2"></td><td colspan="3">2-way ANOVA</td></tr>
<tr><td>Interaction</td><td>Genotype</td><td>Temperature</td></tr>
<tr><td rowspan="2">Supp Fig 2A</td><td></td><td>Kurtosis</td><td>F(1, 8)=2.36<br>P=0.1633</td><td>F(1, 8)=0.31<br>P=0.5901</td><td>F(1, 8)=4.72<br>P=0.0616</td></tr>
<tr><td></td><td>Skewness</td><td>F(1, 8)=0.09<br>P=0.7691</td><td>F(1, 8)=0.44<br>P=0.5270</td><td>F(1, 8)=5.23<br>P=0.0516</td></tr>
<tr><td colspan="3" rowspan="2"></td><td colspan="3">Unpaired Student´s t-test</td></tr>
<tr><td>Df</td><td>t</td><td>P value</td></tr>
<tr><td rowspan="2">Supp Fig 2B</td><td rowspan="2">30°C</td><td>T3</td><td>4</td><td>3.441</td><td>0.0263</td></tr>
<tr><td>T4</td><td>4</td><td>0.4327</td><td>0.6876</td></tr>
<tr><td colspan="3" rowspan="2"></td><td colspan="3">2-way ANOVA</td></tr>
<tr><td>Interaction</td><td>Genotype</td><td>Time</td></tr>
<tr><td rowspan="2">Supp Fig 2C</td><td rowspan="2">30°C</td><td>MAX</td><td>F(11, 44)=3.23<br>P=0.0027</td><td>F(1, 4)=7.26<br>P=0.0544</td><td>F(11, 44)=8.51<br>P<0.0001</td></tr>
<tr><td>MIN</td><td>F(11, 44)=1.36<br>P=0.2275</td><td>F(1, 4)=0.00<br>P=0.9906</td><td>F(11, 44)=3.39<br>P=0.0018</td></tr>
<tr><td colspan="3" rowspan="2"></td><td colspan="3">2-way ANOVA</td></tr>
<tr><td>Interaction</td><td>Genotype</td><td>Treatment</td></tr>
<tr><td rowspan="2">Supp Fig 2E</td><td rowspan="2">30°C</td><td>Kurtosis</td><td>F(1, 4)=0.97<br>P=0.3793</td><td>F(1, 4)=14.23<br>P=0.0196</td><td>F(1, 4)=0.046<br>P=0.8405</td></tr>
<tr><td>Skewness</td><td>F(1, 4)=11.68<br>P=0.0268</td><td>F(1, 4)=0.089<br>P=0.7799</td><td>F(1, 4)=0.56<br>P=0.4964</td></tr>
<tr><td rowspan="2">Supp Fig 2F</td><td rowspan="2">30°C</td><td>Pln/Atp2a2</td><td>F(1, 19)=0.34<br>P=0.5684</td><td>F(1, 19)=0.09<br>P=0.7700</td><td>F(1, 19)=21.86<br>P=0.0002</td></tr>
<tr><td>Dio2</td><td>F(1, 13)=3.37<br>P=0.0894</td><td>F(1, 13)=8.31<br>P=0.0128</td><td>F(1, 13)=4.66<br>P=0.0502</td></tr>
</table>
